# Multifunctional RNA-binding proteins influence mRNA abundance and translational efficiency of distinct sets of target genes

**DOI:** 10.1101/2021.04.13.439465

**Authors:** Valentin Schneider-Lunitz, Jorge Ruiz-Orera, Norbert Hubner, Sebastiaan van Heesch

## Abstract

RNA-binding proteins (RBPs) can regulate more than a single aspect of RNA metabolism. We searched for such previously undiscovered multifunctionality within a set of 143 RBPs, by defining the predictive value of RBP abundance for the transcription and translation levels of known RBP target genes across 80 human hearts. This led us to newly associate 27 RBPs with cardiac translational regulation *in vivo*. Of these, 21 impacted both RNA expression and translation, albeit for virtually independent sets of target genes. We highlight a subset of these, including G3BP1, PUM1, UCHL5, and DDX3X, where dual regulation is achieved through differential affinity for target length, by which separate biological processes are controlled. Similar to the RNA helicase DDX3X, the known splicing factors EFTUD2 and PRPF8 - all identified as multifunctional RBPs by our analysis - selectively influence target translation rates depending on 5’ UTR structure. Our analyses identify dozens of RBPs as being multifunctional and pinpoint potential novel regulators of translation, postulating unanticipated complexity of protein-RNA interactions at consecutive stages of gene expression.

## INTRODUCTION

RNA-protein interactions are crucial for a wide range of processes in multiple subcellular compartments, including RNA transcription, splicing, editing, transport, stability, localization, and translation (Gerstberger et al., 2014). Using state-of-the-art mass spectrometry-based approaches, recent studies have identified up to thousands of potential RNA-binding proteins (RBPs), although for many their precise roles remain unknown (Caudron-Herger et al., 2019; Mallam et al., 2019; Trendel et al., 2019). Whereas RBPs can interact with target RNAs through well-defined protein-RNA binding domains (RBD), uncharacterized RBDs are frequently discovered (Castello et al., 2016; Beckmann et al., 2016) highlighting the complex, diverse, and still largely unknown nature of RNA-protein interactions.

The possession of more than a single RBD could theoretically allow RBPs to be multifunctional, for instance through the separate regulation of different sets of targets (Kong et al., 2019). Multifunctionality may additionally be established through the condition- or cell type-specific expression of RBPs and their interaction partners, or through dynamic shuttling of RBPs between different subcellular compartments, such as the nucleus and cytosol (Dassi, 2017; Díaz-Muñoz & Turner, 2018; Gray et al., 2015; Yugami et al., 2020). Accordingly, out of a large number of RBPs whose subcellular localization was recently evaluated (Van Nostrand et al., 2020), the vast majority could be detected in more than one compartment, albeit of thus far largely unknown biological significance. The potential importance of RBP shuttling was exemplified by the subcellular redistribution of dozens of RBPs upon global induction of mRNA decay (Gilbertson et al., 2018), and compartment-specific changes in RBP interactomes that could be witnessed upon cellular stress (Backlund et al., 2020). These observations indicated that RBPs may dynamically relocate to form a mechanistic bridge between normally compartment-restricted consecutive stages of gene expression regulation: some taking place in the nucleus (e.g., transcription, mRNA processing, splicing), others in the cytosol (e.g., translation, RNA decay).

RBPs are crucial regulators of mRNA translation – the synthesis of protein from a messenger RNA template by ribosomes (Hinnebusch, 2014). In the cytosol, RBPs can act in a general capacity (e.g., mRNA translation initiation or elongation), as well as in more specialized functions where select accessory RBPs or heterogeneous ribosomes facilitate the translation of dedicated subsets of target RNAs (Harvey et al., 2018b; Shi et al., 2017; Simsek et al., 2017). In the nucleus, RBPs can coordinate the process of splicing and translation by modulating different structural and sequence properties of the mRNAs that are exported to the cytoplasm. For instance, highly structured regions in 5’ UTRs can decrease the efficiency of translation initiation at the cost of an overall lower translational output (Lim et al., 2018; Wang et al., 2020; Leppek et al., 2018; Pelletier & Sonenberg, 1985), whereas increased RNA structures in CDS or 3’UTR regions may also enhance transcript stability and hence mRNA half-life, in turn yielding higher protein output over time (Mauger et al., 2019; Cheng et al., 2020). Additionally, alternative UTR usage can expose translated uORFs or miRNA binding sites affecting mRNA translation and/or stability. Other alternative splice events might affect intrinsic properties of an mRNA, such as the ‘swiftness’ of nuclear export (Katahira, 2015), codon usage (Drummond & Wilke, 2008; Qian et al., 2012), affinity to the exon junction complex (EJC) (Nott et al., 2004), or susceptibility to nonsense-mediated decay (Lewis et al., 2003).

Mechanistic insight into the quantitative effects of RBP expression on mRNA translation - i.e., when translation rates of the endogenous RBP targets respond directly to changes in RBP abundance - was recently provided by several studies (Chothani et al., 2019; Luo et al., 2020; Sharma et al., 2021). These studies used this quantitative relationship to assign novel functions to known RBPs and to investigate the kinetics through which RBPs regulate their targets. For instance, Chothani et al. used a computational correlation approach to define the frequency with which RBP levels correlate with target translation rates. This helped to pinpoint key RBP network hubs that were crucial for translational regulation during cardiac fibrosis *in vitro* and *in vivo* (Chothani et al., 2019). In a separate study, Luo et al. used luciferase-based 3’ UTR tethered function assays for 690 RBPs, identifying 50 RBPs whose expression induced significant positive or negative effects on mRNA stability and/or translation. This resulted in the novel characterization of the stress granule RBP UBAP2L as a ribosome-associated RBP (Luo et al., 2020). Lastly, Sharma et al. developed a methodology to investigate RNA-protein kinetics in a time-resolved manner for the RBP DAZL (Sharma et al., 2021). This approach helped establish a quantitative relationship that precisely explained the effect of DAZL on mRNA levels and ribosome association, which correlated with the cumulative probability of DAZL binding within clusters of proximal 3’ UTR binding sites.

In the current study, we describe the computational identification of putative new, and sometimes dual functions of RBPs in the regulation of mRNA translation levels. We integrate crosslinking immunoprecipitation (CLIP)-derived mRNA targets for 143 RBPs (Van Nostrand et al., 2020; Maatz et al., 2014) with the transcriptomes and translatomes of 80 human hearts (Van Heesch et al., 2019) – a tissue where translational control is known to play a central role in gene expression regulation (Chothani et al., 2019; Van Heesch et al., 2019; Schafer et al., 2015b). We show that the expression levels of many, but not all, investigated RBPs indeed correlate with target mRNA abundance and/or translational efficiency *in vivo*. For a subset of 21 RBPs (including proteins with diverse previously described roles, such as the endoribonuclease G3BP1, the helicase DDX3X, the protease PUM1 and the deubiquitinase UCHL5), we could independently assign dose-dependent effects to both mRNA levels and translational efficiencies of largely distinct sets of target genes each involved in unrelated biological processes. Mechanistically, these target genes also appeared to be regulated independently, facilitated by a differential affinity for coding sequence length or 5’ UTR structure.

Our results show that RBPs with more than a single role are likely to be more prevalent than currently anticipated. We postulate that multifunctional RBPs may use their functional plasticity in a condition- or compartment-specific manner to regulate gene expression at multiple levels, for separately defined sets of target genes.

## RESULTS

### RNA-binding protein abundance determines the efficiency of target gene translation

With the aim to define which RBPs are capable of influencing more than a single stage of gene expression control, we first determined whether RBP abundance can have predictive value for the extent of target gene regulation in the human heart. To this extent, we compiled protein-RNA interactions for 143 cardiac-expressed RBPs, consisting of the muscle-specific RBM20 (Maatz et al., 2014) and 142 ubiquitously expressed RBPs previously characterized in depth as part of ENCODE (Van Nostrand et al., 2020) (see Methods, **Figure S1A** and **Table S1**). We then correlated the abundance of these 143 RBPs with mRNA levels and translational efficiencies (TE) of 11,387 cardiac-expressed genes across 80 human hearts - the largest set of matched human tissue transcriptomes and translatomes that is currently available (Van Heesch et al., 2019). This revealed a clear quantitative relationship between RBP expression levels and the extent of target gene expression control (e.g., translation rates: **Figure 1A** and **Figure S1B**).

**Figure 1.**
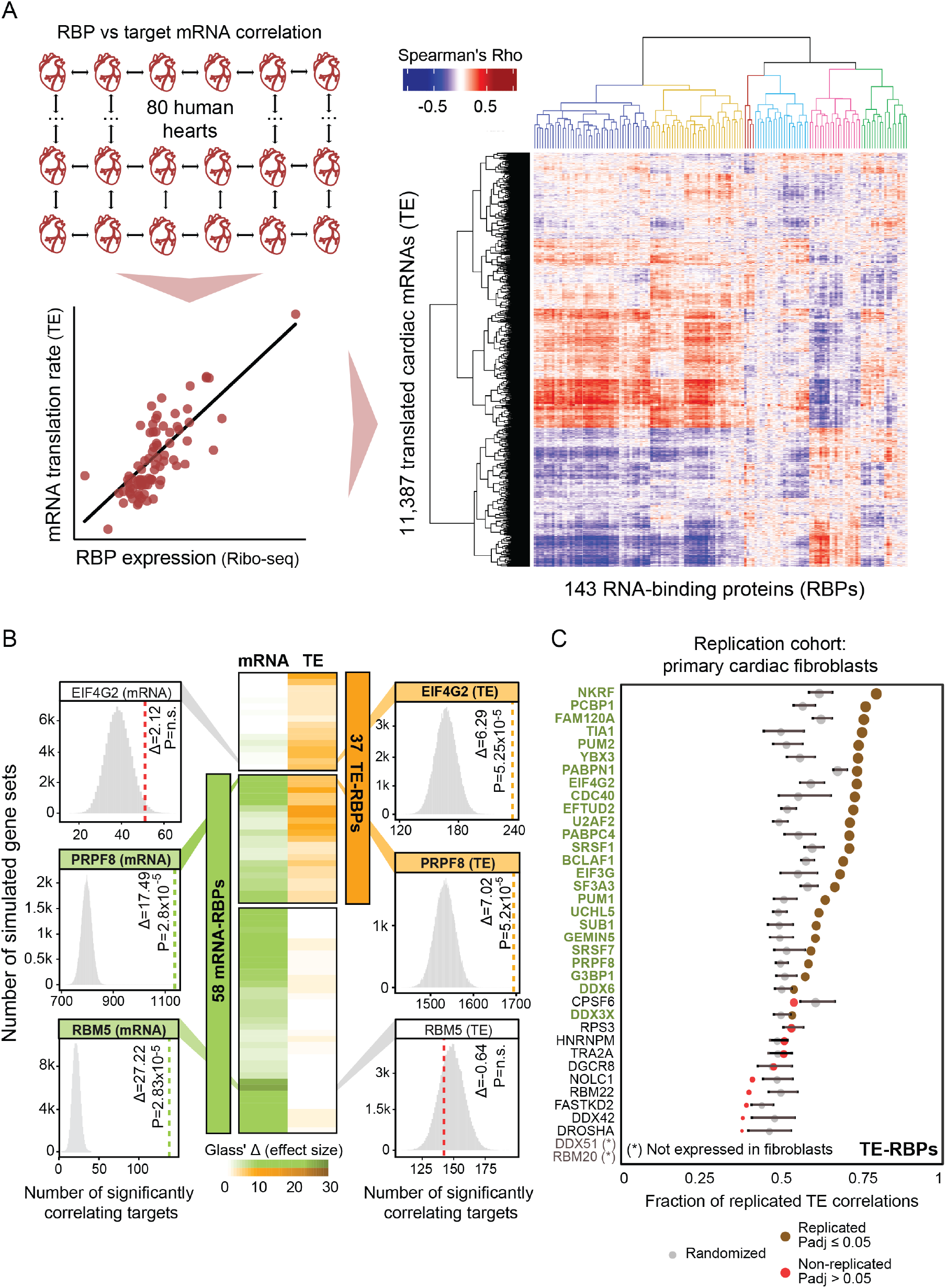
RNA-binding protein abundance predicts target translational regulation. **(A)** Schematic of the RBP-target correlation approach. Using the quantified Ribo-seq and RNA-seq data from 80 hearts, pairwise RBP versus target mRNA abundance or translational efficiency correlations were calculated. A heatmap with hierarchically clustered translational efficiency Spearman’s Rho correlations of RBPs and translated mRNAs in the human heart are shown. Six clusters of coregulated RBPs are highlighted (See also **Table S1**). **(B)** Heatmap with Glass’ Δ scores that quantify the effect size of the witnessed significance of associations between RBPs and target gene mRNA abundance and TE. Only significant RBPs are shown: 37 TE-RBPs (orange) and 58 mRNA-RBPs (green). For three selected RBPs (one per category), histograms illustrate the significance of the calculated associations. **(C)** Dot plot displaying the fraction of translational efficiency RBP-target correlations that can be replicated in an independent set of primary cardiac fibroblasts (Chothani et al., 2019). For each RBP, the significance of the replication was evaluated by comparing the replicated fraction between observed and randomized sets and it is represented as a brown (significant) or red (non-significant) dot. The size of the dots indicates the strength of significance (-log10 (p_adj_)) and grey dots correspond to the fraction of replicated correlations in randomized sets. Error bars indicate mean values with standard deviation (SD).

Next, we calculated the frequency with which mRNA levels and translational efficiencies of the CLIP-derived target genes correlated significantly with the abundance of each RBP. For this, we statistically evaluated associations through the sampling of 100,000 matched sets of simulated target genes, yielding 58 RBPs with significant associations (empirical p_adj_ ≤ 0.05) with target mRNA abundance (hereafter denoted as “mRNA-RBPs” that regulate “mRNA targets”) and 37 RBPs with significant associations with target translational efficiencies (“TE-RBPs” regulating “TE targets”) (**Figure 1B** and **Figure S1C**). These, for instance, included the candidate tumor suppressor and ubiquitous splicing regulator RBM5 (Sutherland et al., 2010), which we identified as a cardiac mRNA-RBP influencing mRNA abundance of at least 138 correlating targets (p_adj_ = 2.83 × 10^−5^; Glass’ Δ = 27.2). Reassuringly, our strategy validated known TE-RBPs such as the eukaryotic translation initiation factor EIF4G2, whose expression dynamics could be associated with target gene translational efficiencies of at least 235 correlating targets expressed in the human heart (p_adj_ = 5.26 × 10^−5^; Glass’ Δ = 6.3) (Liberman et al., 2009; Weber R, Kleemann L, Hirschberg I, Chung M, Valkov E, 2021) (**Figure 1B**). Importantly, we could replicate our calculations for 25 out of 37 depicted TE-RBPs in an independent, though smaller cohort of primary cardiac fibroblast translatomes (n = 20) (Chothani et al., 2019) - a system previously explored to identify RBPs with key roles in cardiac fibrosis (**Figure 1C**).

### Positive and negative control of translation by known and unknown factors

We detected 27 (out of 37) TE-RBPs without prior evidence of regulating mRNA translation, including 4 RBPs with no function assigned to their RNA binding ability at all (NKRF, FAM120A, SUB1 and UCHL5) (**Figure S2A**). In order to define how these RBPs interact with existing RBP networks to regulate target gene translation in a coordinated manner, we hierarchically clustered the correlation coefficients of all 37 TE-RBPs and their CLIP targets. This primarily divided the matrix in two distinct groups of TE-RBPs with marked opposite effects on target TE, indicative of competition and/or cooperation between subsets of RBPs (**Figure 2A**).

**Figure 2.**
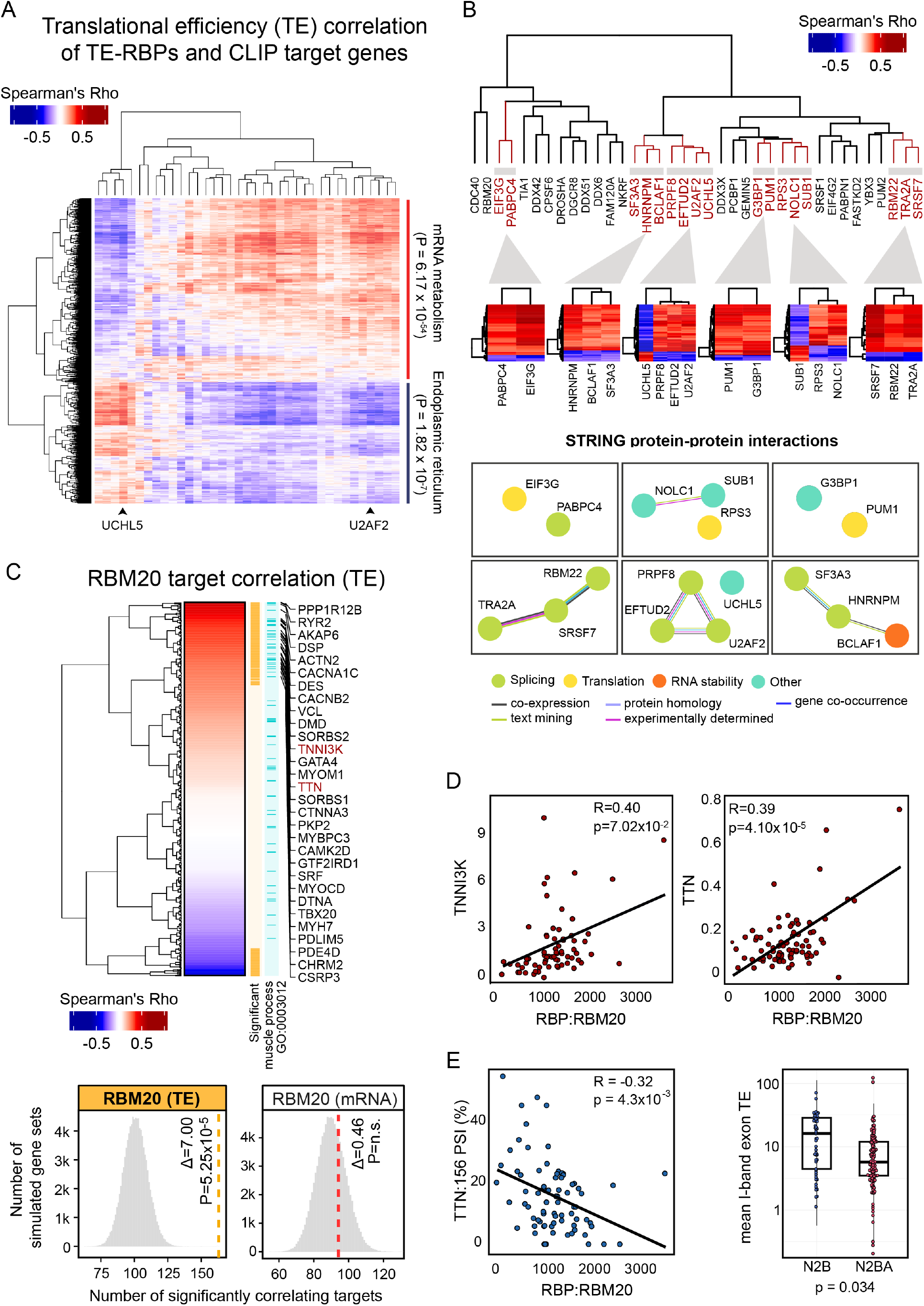
CLIP analysis identifies coregulated *in vivo* targets and novel master translational regulators in the human heart. **(A)** Heatmap with hierarchically clustered translational efficiency correlations of TE-RBPs and predicted target genes in the human heart is represented. The clustering separates two groups with opposite effects on TE whose targets are enriched in mRNA metabolism (p_adj_ = 6.17 × 10^−54^) and endoplasmic reticulum (p_adj_ ≤ = 1.82 × 10^−7^) GO terms respectively. **(B)** Dendrogram with hierarchically clustered TE-RBPs based on pairwise RBP-RBP overlaps. Shared target genes of all paired RBPs were included for clustering. Bottom heatmaps with translational efficiency correlations of selected RBP clusters and shared significant targets. These plots illustrate distinct cooperative and competitive RBP-target regulation modes. STRING protein-protein interaction networks (Szklarczyk et al., 2019) from selected RBP clusters reveal functional association of coregulated RBPs. Colours in edges and nodes indicate the sources of STRING evidence and known RBP functions. **(C)** Heatmap with hierarchically clustered Spearman’s Rho correlation scores of RBM20 and the translational efficiency of the predicted target genes. Significant correlating targets (n = 163, p_adj_ ≤ 0.05) and targets involved in muscle process (GO: 0003012) are highlighted in orange and light blue colours respectively. A list of sarcomere gene targets positively correlating with RBM20 is displayed. Selected bottom histograms illustrate the significance of RBM20 with correlating TE targets and the absence of significance with correlating mRNA targets. **(D)** Scatter plots representing the correlation of translational efficiencies between RBM20 and two sarcomere genes, *TNNI3K* and *TTN*. Score and level of significance of the two Spearman’s correlations are displayed. **(E)** Left: Scatter plot showing correlation between RBM20 and *TTN* exon:156 percent spliced in (PSI). Score and level of significance of the two Spearman’s correlations are displayed. Right: Box plot comparing average *TTN* I-Band isoform-specific TEs. N2B (ENST00000460472) has a significantly higher TE than N2BA (ENST00000591111) (p-value = 0.034).

Interestingly, the presented method joins known and novel TE-RBPs with opposing or concordant directionality of regulation on shared targets (**Figure 2B**). For instance, depending on the shared target gene bound by the splicing factor U2AF2 and the protease UCHL5, completely opposite effects on TE could be observed and independently replicated (**Figure 2C, Figure S2B** and **S2C**). Although these shared modes of target regulation were in part concordant with protein-protein interactions annotated in the STRING database (Szklarczyk et al., 2019), a subset of coregulatory “RBP hubs” contained proteins with previously unknown functional similarities (e.g., UCHL5 and U2AF2).

Amongst the 27 potentially new TE-RBPs was the muscle-specific and dilated cardiomyopathy-associated splicing regulator RBM20, whose expression correlated particularly well with the TE of 163 experimentally validated target genes (out of 561 total targets; Glass’ Δ = 7.0; **Figure 2C**). Importantly, RBM20 levels specifically influenced TE and had no impact on overall mRNA abundance or stability. Most RBM20 targets, including the sarcomere genes *TTN* and *TNNI3K*, correlated positively (i.e., higher RBM20 expression associates with increased target gene TE; **Figure 2D**) and especially those positively correlating targets showed strong enrichment for muscle function processes (GO:0003012, p_adj_ ≤ 5.97 × 10^−16^) (**Figure 2C**). To investigate a possible connection between RBM20-mediated mRNA splicing and the subsequent efficiency of mRNA translation, we evaluated whether splicing rates of known target exons correlated directly with TE. For 66 out of 163 (± 40%) translationally regulated RBM20 target genes, the extent of alternative splicing (PSI; see methods) indeed correlated with RBM20 abundance. A clear example is the exon inclusion measured across the *TTN* I-band, whose exons are only included in the longer *TTN* N2BA isoform. These I-band exons specifically drive the negative correlation of RBM20 expression with overall *TTN* TE, indicating that their inclusion reduces the efficiency with which *TTN* can be translated. We had previously observed that *TTN* translation rates are strongly isoform-dependent (van Heesch et al, 2019) and can now mechanistically connect this to splicing control by RBM20 (**Figure 2E**), a consequence that seems generalizable for more muscle-specific RBM20 targets, including other sarcomere components (**Figure 2D** and **2E**). Although the precise mechanism through which RBM20 influences TE remains unknown, this RBP may omit the inclusion of exons with inefficient codon translation rates or exons that impact the stability or structure of the transcript, both of which influence protein synthesis rates (Nott et al., 2004; Zhou et al., 2016).

### Multifunctional RBPs including DDX3X and G3BP1 regulate mRNA abundance and translational efficiency of independent sets of target genes

Among the 74 RBPs that correlated significantly with target gene TE (37 TE-RBPs) or mRNA abundance (58 mRNA-RBPs), a subset of 21 RBPs could be associated independently with both of these molecular traits (**Figure 1B** and **3A**). To investigate whether the association with mRNA abundance and TE was interrelated (and hence successive, i.e., higher target expression drives increased TE), we examined the sets of genes that correlated significantly with either trait. This revealed a very limited overlap between correlating target genes for all 21 RBPs (16.71 ± 8.19%, **Figure 3A** and **Table S2**), which decreased further for the most strongly correlating targets (1.09 ± 1.49%; r > 0.5; **Figure S3A**). To substantiate this observation, we compared the trait-specific strength of the effect sizes and found no relation between the correlations as independently witnessed for both traits, confirming the absence of a carryover effect (**Figure 3B** and **Figure S3A**). This led us to denote these RBPs as “multifunctional RBPs” - context-specific RBPs whose functional outcome depends on the set of mRNAs it targets. A key example appears to be the multifunctional RBP DDX3X (Mo et al., 2021; Soto-Rifo et al., 2012), whose abundance correlates significantly with the mRNA levels of 339 target genes (p_adj_ = 2.83 × 10^−5^; Glass’ Δ = 6.9) and the translational efficiency of 730 target genes (p_adj_ = 5.25 × 10^−5^; Glass’ Δ= 11.89), of which only 43 targets overlap between both sets (**Figure 3A** and **3C**). The consequences of DDX3X binding for mRNA abundance (positive correlation) or TE (negative correlation) are opposite, though this is not the case for all multifunctional RBPs (**Figure S3B**). Three other multifunctional RBPs similarly act as repressors of translation whilst having a positive effect on mRNA abundance (DDX6, NKRF, GEMIN5), one RBP shows the exact opposite behavior (FAM120A), and all others have concordant roles at both layers of control (e.g., TRA2A, FASTKD2, SRSF1).

**Figure 3.**
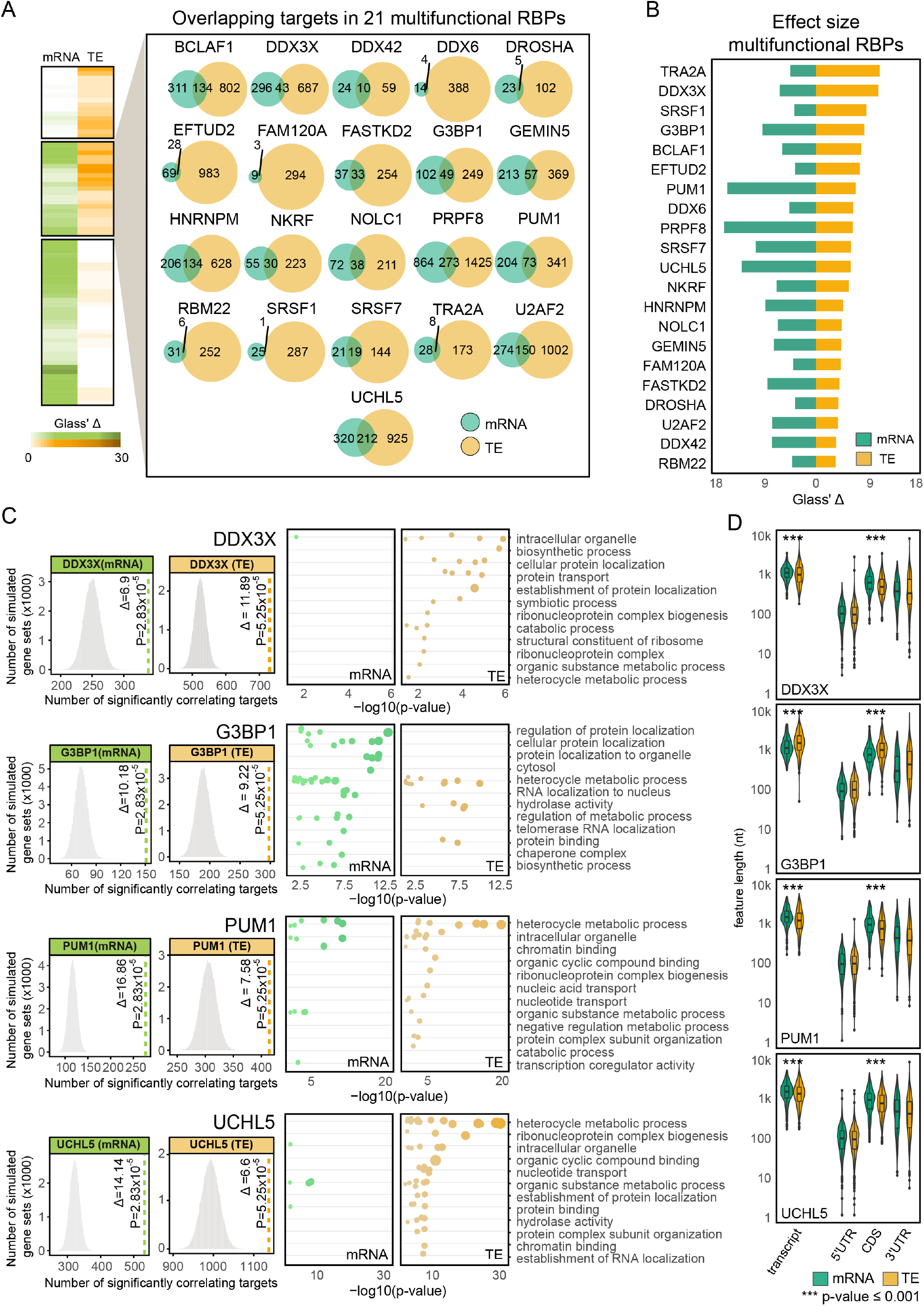
Multifunctional RBPs regulate translation of distinct sets of target genes. **(A)** Heatmap with Glass’ Δ scores quantifying the effect size of the witness effects for mRNA and TE correlations. Both effect sizes are significant for a highlighted set of 21 multifunctional RBPs. For this set of RBPs, individual Venn Diagrams representing the overlap in the total number of mRNA and TE targets are displayed. **(B)** Bar plot quantifying the magnitude of mRNA and TE effect size (Glass’ Δ scores) for multifunctional RBPs. RBP effect sizes are largely independent of the mode of regulation. **(C)** Selected histograms and dot plots illustrating the significance of RBP-target correlations and the enrichment of GO terms for the targets bound by 4 multifunctional RBPs: DDX3X, G3BP1, PUM1, and UCHL5. For each RBP, the 12 most significant parental GO terms are displayed. For three of the RBPs, mRNA and TE targets exhibit different enrichment of significant GO terms. **(D)** Box plots with transcript, 5’ UTR, CDS, and 3’ UTR sequence lengths in nucleotides for mRNA and TE targets corresponding to the four selected multifunctional RBPs in (C). A total of 11 multifunctional RBPs bind targets with significantly different CDS lengths. See also **Figure S3**.

Of note, the TE and mRNA target genes of multifunctional RBPs can be enriched for completely separate biological processes, indicating that duality can contribute to independent biological outcomes. For instance, correlating DDX3X and UCHL5 TE targets code for proteins involved in RNA splicing (GO:0008380, p_adj_ = 7.70 × 10^−30^), while their mRNA targets did not show any clear functional enrichment (**Figure 3C**). For G3BP1, mRNA targets code for proteins involved in localization to nuclear body (GO:1903405, p_adj_ = 5.13 × 10^−12^), whereas this is not the case for translationally regulated targets, which are enriched for RNA splicing (GO:0008380, p_adj_ = 1.61 × 10^−10^). Such biological discrepancies are not always present: independent of the mode of regulation, both types of PUM1 targets (TE or mRNA) appear to code for proteins involved in mRNA processing (GO:0006397, TE p_adj_ = 2.64 × 10^−22^, mRNA p_adj_ = 6.40 × 10^−13^).

### Differential affinity of multifunctional RBPs for CDS lengths and 5’ UTR structures

Dual- or multiprotein functionality can be achieved through context-specific differences in subcellular localization (Buchan, 2014), interaction partners (Cirillo et al., 2020; Yang et al., 2019), or the presence of multiple RNA-binding domains (Müller-McNicoll & Neugebauer, 2013) - all of which can fine tune or restrict the subset of recognized target genes. Based on published immunofluorescence imaging-based evidence of subcellular RBP localization (Van Nostrand et al., 2020), 13 out of 21 multifunctional RBPs indeed localized equally well to both nucleus and cytosol, suggesting functionality in both compartments (**Table S3**).

We additionally explored the relative position of CLIP binding sites in target genes (i.e., the position of binding within the mRNA: 5’ or 3’ UTR, CDS, or intronic). Most of the RBPs (including e.g., DDX3X) showed no marked difference in binding positioning (**Figure S3B**). For DDX3X and 10 other RBPs, we did notice a significant change in target transcript length, mostly explained by differences in target CDS length, which slightly increased or decreased between TE and mRNA targets (**Figure 3D** and **Figure S3C**). The most significant changes in CDS length were seen for GEMIN5 (decrease for TE targets; 2,226nt vs. 1,519nt; p_adj_ = 3.66 × 10^−9^), PRPF8 (decrease for TE targets; 2,243nt vs. 2,076nt; p_adj_ = 1.03 × 10^−8^), DDX3X (decrease for TE targets; 1,659nt vs 1,376nt; p_adj_ = 2.81 × 10^−7^) and G3BP1 (increase for TE targets; 1,985 vs. 2,798nt; p_adj_ = 6.11 × 10^−8^).

Our attention was drawn to the DEAD-box helicase DDX3X, which regulates translation initiation by interacting with, and subsequently disentangling, highly structured RNA sequences (Guenther et al., 2018; Soto-Rifo et al., 2012; Sen et al., 2015; Calviello et al., 2020). DDX3X binds 5’ UTRs and the small ribosomal unit to facilitate the translation of a subset of mRNAs with long and structured leader sequences (Calviello et al., 2020). In order to define if additional RBPs may be required for, or involved in, translation initiation at targets with highly structured 5’ UTRs, we looked into the 5’ UTR minimum free energy (MFE, length normalized) of TE and mRNA target genes of all multifunctional RBPs. We observed that between the positively and negatively correlating target translational efficiencies of 17 out of 21 multifunctional RBPs, 5’ UTR sequences differed in structural composition (**Figure 4A**). In contrast, there were poor to almost no differences for the significantly correlating mRNA targets of the same RBPs. Strikingly, three RBPs exhibited by far the strongest MFE differences between positively and negatively correlating targets: next to DDX3X (p_adj_ = 9.47 × 10^−47^), those were the core spliceosome factors PRPF8 (p_adj_ = 2.70 × 10^−29^) and EFTUD2 (p_adj_ = 1.69 × 10^−30^) (**Figure 4A** and **4B**). The targets shared between these RBPs displayed similar directions of correlation with the three RBPs (**Figure 4C**) and code for proteins involved in mRNA stabilization (GO:0048255, p_adj_ = 3.09 × 10^−4^), suggesting that these three RBPs might rely on a similar mechanism for translational regulation that is dependent on the 5’ UTR structures of shared targets.

**Figure 4.**
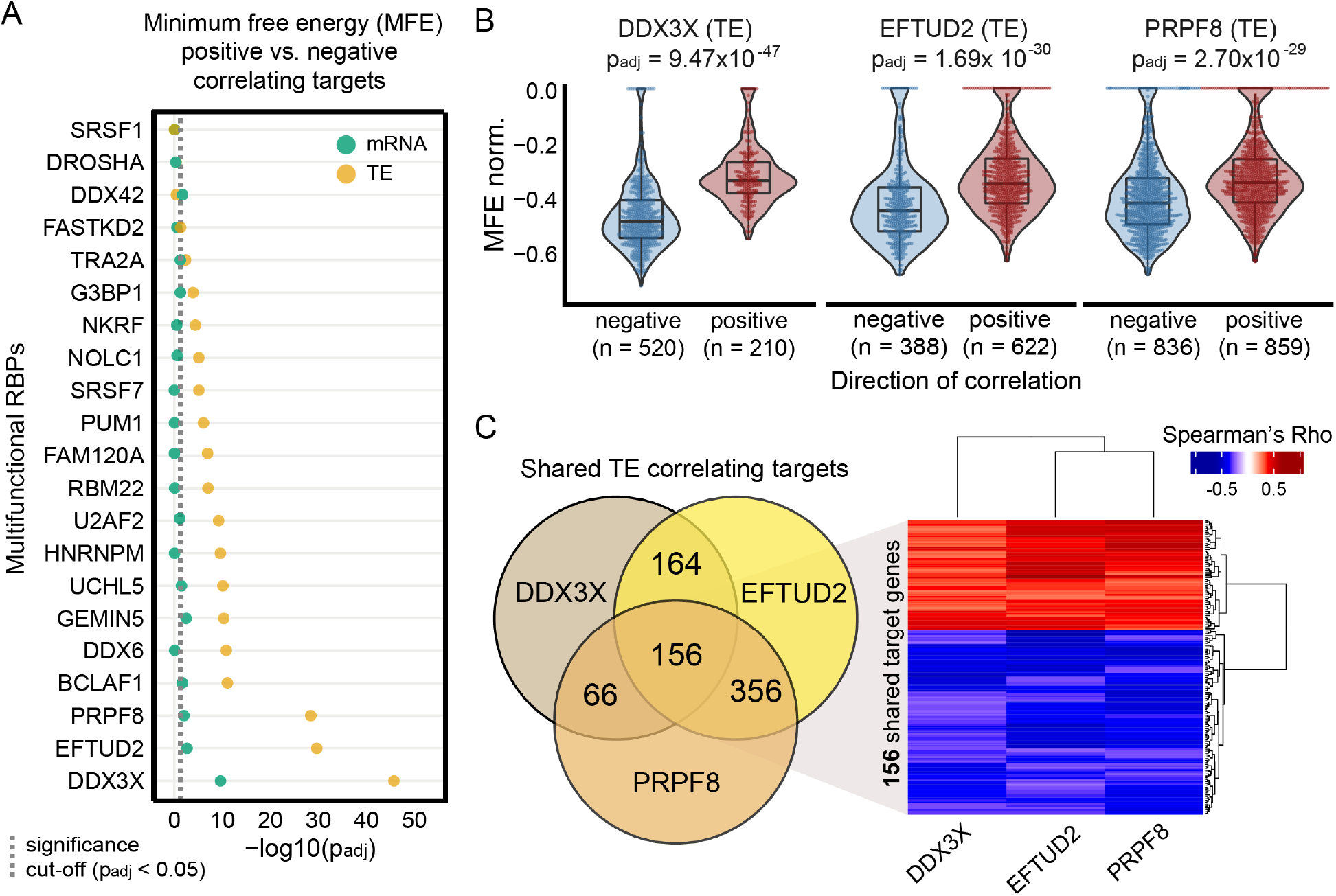
Differential affinity of multifunctional RBPs for 5’ UTR structures often drives opposite quantitative TE effects. **(A)** Dot plot displaying the significance of the differences in 5’ UTR minimum free energy (MFE, normalized by length) between target genes that correlate positively or negatively with each multifunctional RBP. Significance values are calculated separately for mRNA (green) and TE (brown) targets. Adjusted p-values are shown on a -log10 scale and calculated using the Wilcoxon Rank Sum test and only 5’ UTR sequences with a minimum length of 20 nucleotides were evaluated. A dashed vertical line indicates the minimum adjusted p-value to consider the differences in MFE as significant (p_adj_ < 0.05). **(B)** Box and violin plots with length normalized MFE scores for positively and negatively correlated TE targets corresponding to the three selected multifunctional RBPs with the highest significance in Figure 4A (DDX3X, EFTUD2, PRPF8). **(C)** Three-way Venn Diagram representing the overlap in the number of TE targets for the three selected RBPs. The heatmap represents TE correlations of 156 shared target genes for the three cases.

Collectively, our results point to an intricate relation between the direction of translation regulation and target 5’ UTR structure, with the TE rates of certain targets being positively or negatively influenced by multifunctional RBP binding depending on the complexity of target 5’ UTR sequences.

## DISCUSSION

Increasing evidence suggests that RBPs can act as multifunctional gene expression regulators (Backlund et al., 2020; De Gaudenzi et al., 2016). Here, we built an in-silico method for the large-scale analysis of RBP-driven regulation using correlation as a proxy for mRNA abundance and translational efficiency (TE) of target genes across 80 human heart samples. Our approach underscores the functional importance of RBP expression fluctuations in the control of gene expression, a mechanistic feature recently highlighted by others *in vitro* and *in vivo* to assign new unknown roles for RBPs (Chothani et al., 2019; Luo et al., 2020; Sharma et al., 2021). We exploited the quantitative effect of RBPs on known target genes to implicate 74 RBPs in the regulation of mRNA abundance and translation.

We discovered 27 RBPs with previously unknown roles in translation, some of which have well-characterized functions in other biological processes, including mRNA splicing. Previous work revealed a handful of splicing factors that can independently mediate post-splicing activities, such as mRNA translation (Chen et al., 2019; Maslon et al., 2014; Kim et al., 2014). Instead, alternative splicing mediated by splicing factors can cause downstream effects on the translation of the spliced target genes depending on the qualitative decision of which mRNA isoforms are being produced (Lim et al., 2018; Wang et al., 2020; Leppek et al., 2018; Pelletier & Sonenberg, 1985; Mauger et al., 2019; Cheng et al., 2020; Katahira, 2015; Drummond & Wilke, 2008; Qian et al., 2012; Nott et al., 2004; Lewis et al., 2003). Remarkably, the high fraction of splicing factors that we find to influence translation suggests previously unanticipated roles for many more splicing regulators in this process. A prominent example is the muscle-specific and cardiac disease-relevant splicing regulator RBM20.

For the first time, we demonstrate that RBM20 expression correlates positively with the TE of many sarcomere genes, suggesting that the nuclear splicing control can impact cytoplasmic protein synthesis. Future functional studies should determine the specific mode of action of the splicing factors involved in translation and whether any isoform-specific characteristics contribute to the observed differences in TE.

Besides the discovery of potential novel functions for a subset of RBPs, we provide evidence that 21 RBPs can modulate both target mRNA abundance and TE - a class of RBPs that we classify as “multifunctional RBPs”. Interestingly, multifunctional RBPs appear to be involved in the regulation of mRNA abundance and TE of distinct groups of target genes. These target genes can be concordantly or discordantly regulated on either layer of gene expression control. We discovered that the specific affinity of several RBPs to structural properties of mRNAs, such as protein-coding sequence length, UTR length or RNA secondary structure, contribute separately to the observed independent effects on mRNA abundance or translation. Hence, further experimental validation should be required in order to confirm the quantitative effect of each individual multifunctional RBP and identify the precise mechanisms behind this dual functionality. In support of our findings, a recent study (Sharma et al., 2021) has inspected the connection between the binding kinetics of the RBP DAZL and their effect on mRNA abundance and translation of specific sets of target genes, identifying several 3’ UTR features – UTR length, presence of binding clusters, distance to the polyadenylation site – that are linked to the trait-specific regulation of different groups of targets. In addition, the usage of different ribosome binding domains, the recognition of alternative RBP motifs and the presence of binding sites located in different gene regions (i.e., UTRs, CDS, or introns) can also be indicative of RBP multi-functionality (Ray et al., 2013). Nevertheless, we found these mRNA characteristics to remain largely unchanged for the targets of multifunctional RBPs identified in this study.

For a subgroup of multifunctional RBPs, we noticed that the targets regulated on the transcriptional or translational level represent functionally different gene classes. Indeed, the observed biological diversity in our study seems to match the condition-specific regulatory complexity that needs to be achieved by a single RBP. The limitations of the present study for assigning functionality naturally include the use of eCLIP experiments from two cell lines unrelated to heart. Our results would benefit from large-scale human heart-specific CLIP experiments that better complement the tissue-specificity of the cardiac expressed genes. Nevertheless, all discovered multifunctional RBPs are ubiquitously expressed across human tissues, suggesting that these proteins are major players in RNA transcriptional and translational regulation, contributing to both global and cell-line or tissue-specific manifestations. For instance, this appears to be the case for G3BP1 - a known multi-functional RBP that can selectively compartmentalize specific sets of mRNAs to stress granules, in order to reprogram mRNA translation under certain global stress conditions (Matsuki et al., 2013; Ying & Khaperskyy, 2021; Sahoo et al., 2018), as well as tissue-specific contexts such as atrial fibrillation in heart (Dong et al., 2019). Additionally, G3BP1 plays an important role in DNA/RNA unwinding (Costa et al., 1999) and binds to specific RNA stem-loop structures to trigger mRNA degradation (Fischer et al., 2020), which is essential for maternal mRNA clearance (Laver et al., 2020)). Another example is DDX3X, a DEAD box helicase which can respond to stressors (e.g., viral infections (Yedavalli et al., 2004)) by switching subcellular compartments. DDX3X is involved in multiple processes required for RNA metabolism (Mo et al., 2021; Soto-Rifo et al., 2012), for which it uses its capacity to unwind complex and structured 5’ UTRs to promote translation initiation at selected subsets of mRNAs (Soto-Rifo et al., 2012; Shen & Pelletier, 2020; Calviello et al., 2020; Guenther et al., 2018). However, there is ongoing debate as to the precise roles of DDX3X and the mechanisms through which it regulates RNA metabolism (Shen & Pelletier, 2020), as it can act both as a repressor or activator of translation (Hilliker et al., 2011).

Our work demonstrates that multifunctional RBPs can act as either positive or negative regulators of translation based on the structural complexity of target 5’ UTR sequences. Unexpectedly, our results show that increased levels of DDX3X correlate with a lower TE for targets with highly structured 5’ UTRs. This contradicts recent *in vitro* reports where DDX3X knockdown in human cells (Calviello et al., 2020) resulted in translational repression of mRNAs with structured 5’ UTRs. Next to DDX3X, the strongest impact of the 5’ UTR on translational output is observed for EFTUD2 and PRPF8, which display patterns of regulation highly similar to DDX3X, suggesting an analogous mode of action in the control of target translation rates. Surprisingly, EFTUD2 and PRPF8 are splicing factors which are part of the central component of the U5 snRNP spliceosome (Malinová et al., 2017) and had not been implicated in translation before. That the strongest impact of the 5’ UTR on translational output is observed for these two RBPs, suggests a mode of action in the control of target translation rates that is analogous, or closely related to, DDX3X functioning. It would be most probable that these splicing factors define the exon structure of target gene 5’ UTRs, though we have not been able to statistically verify that in our cohort of 80 human heart translatomes. An alternative explanation could be the direct involvement of both RBPs in translation, though only PRPF8 has been observed to shuttle between nucleus and cytosol (Van Nostrand et al, 2020). However, the conserved GTPase EFTUD2 has sequence similarity to the translation elongation factor EF-2 (Fabrizio et al, 1997), which may provide hints toward its capacity to influence translation. Although both ancient paralogs (EF-2 and EFTUD2) may have evolved and diversified to complement each other, further experimental validation is needed to define how EFTUD2 co-defines 5’ UTR structure and thereby translational output.

Whereas dual functionality of extensively studied and characterized RBPs such as DDX3X and G3BP1 had (to a certain extent) been described previously, for a selection of other RBPs our results provide initial observations of dual functionality. For instance, UCHL5 (also known as UCH37) is a protease with RNA binding capacity that may be part of the INO80 chromatin remodeling complex (Yao et al., 2008), though its role within RNA metabolism is yet unknown. We establish a quantitative relationship between UCHL5 expression and variability in mRNA abundance of genes involved in chromatin organization, as well as with changes in TE of genes involved in RNA splicing. Although UCHL5 shares target genes with core splicing factors (U2AF2, EFTUD2 and PRPF8), its effect on the TE of targets shared with these three splicing factors is completely opposite, suggesting contrasting regulatory behavior, and possibly competition.

Very little is known about the molecular processes that control RBP multi-functionality, although some possible mechanisms have been recently investigated, including the formation of heterogeneous RBP complexes (Copsey et al., 2017; Damianov et al., 2016), switches from monomers to multimers in a concentration-dependent manner (Kim & Myong, 2016), and changes in subcellular localization (Chen et al., 2019; Burgess et al., 2011). In our current study, the potential mechanisms behind the observed dual functionality could not be explained in a uniform way: there appears to be no ‘one size fits all’ scenario. It is very probable that RBP multi-functionality is achieved by specific combinations of individual RBP and target features, whose precise dissection requires experimental follow-up into each individual multifunctional RBP. An RBP may bind distinct sets of RNA within the nucleus, though, for a subset of targets, the consequences of binding may only become apparent at a later stage of gene expression (e.g., a change in transcript isoform production that is accompanied by a downstream effect on TE). Alternatively, multifunctional RBPs may physically take part in multiple stages of gene expression by adapting subcellular localizations. For instance, HNRNPM (one of the core splicing ribonucleoproteins that we found to influence both target gene mRNA abundance and TE) localizes to nucleus (Harvey et al., 2018a) but can be exported to the cytosol to induce cap-independent translation upon hypoxia (Chen et al., 2019). Another example, DDX3X, shuttles between nucleus and cytosol (Yedavalli et al., 2004). It remains to be established if RBPs with shared targets bind these targets simultaneously, or if there is a sequential cross-talk of RBPs and other proteins in the control of target expression. RBP abundance may also respond to target availability and not vice versa, possibly explaining why we find many splicing regulators to rank highly within the RBP hierarchy.

In conclusion, our results illustrate unanticipated complexity in RBP-RNA interactions at multiple consecutive levels of gene expression. This warrants future in-depth experimental research into the identified RBPs in human cardiac biology. Understanding how RBPs cooperate, communicate, interact and compete across subcellular compartments and in response to changing conditions will be essential to fully comprehend the quantitative nature of the regulatory principles that underlie mRNA metabolism.

## DATA AVAILABILITY

This study includes no data deposited in external repositories. Accession numbers for all data used in this study have been reported within the Materials and Methods section with reference to the studies that generated and published the data. All scripts generated for the analyses are available via the GitHub development platform at: https://github.com/vschnei/Dual-function_RBP_manuscript_analysis

## ACKNOWLEDGEMENTS

We would like to thank Prof. Dr. Markus Landthaler for his helpful feedback and discussions.

## AUTHOR CONTRIBUTIONS

Conceptualization, V.S-L., N.H. and S.v.H.; Methodology, V.S-L., J. R-O. and S.v.H.; Software, V.S-L. and J. R-O.; Formal Analysis, V.S-L.; Investigation, V.S-L., J. R-O. and S.v.H.; Writing – Original Draft, V.S-L., J. R-O. and S.v.H.; Writing - Review & Editing, all authors; Visualization, V.S-L., J. R-O. and S.v.H.; Funding Acquisition, N.H.; Supervision, S.v.H. and N.H.

## MATERIALS AND METHODS

### Ribosome profiling and RNA sequencing data analysis

We re-analyzed ribosome profiling (Ribo-seq) and matched RNA-seq datasets from 80 human hearts that we generated and published previously (EGA accession code: EGAS00001003263) (Van Heesch et al., 2019). In short, Ribo-seq reads were clipped for residual adapters using FASTX toolkit (Hannon, 2010). Reads mapping to the mitochondrial RNA, ribosomal RNA and tRNA sequences were removed from downstream analysis. Full length paired mRNA-seq reads (2 × 101nt) were trimmed to 29-mers (average length of Ribo-seq reads) so as to establish a comparable analysis of both Ribo-seq and mRNA-seq datasets and avoid any mapping or quantification bias due to different read length or filtering. Next, Ribo-seq and trimmed mRNA-seq reads were mapped to the human reference genome (GRCh38, Ensembl v87) using STAR v2.5.2b (Dobin et al., 2013) with maximum of 2 mismatches and -seedSearchStartLmaxOverLread = 0.5. Quantification of gene expression was performed by counting reads mapping to coding sequence (CDS) regions of annotated protein-coding genes, using HTSeq v0.9.1 (Anders et al., 2015). Gene counts were normalized by estimating the size factors simultaneously on Ribo-seq and RNA-seq datasets using DESeq2 v1.12.4 (Love et al., 2014). This joint normalization is required to compare both measures of gene expression (Zhong et al., 2017). Translational efficiency (TE) was calculated on the Ribo-seq against RNA-seq ratio for each individual gene and sample, as described previously (Van Heesch et al., 2019).

### Identification of RBP targets from published eCLIP and HITS-CLIP data

Processed eCLIP data of 150 RBPs were obtained from ENCODE (Davis et al., 2018) for HepG2 (n = 103) and K562 (n = 120) cell lines. Datasets consisted of BED files containing eCLIP peaks and BAM files containing reads mapped to the human genome (GRCh38.p10/hg38). The identification of robust eCLIP peaks across replicates and cell lines was performed as suggested by Van Nostrand and colleagues (Van Nostrand et al., 2020). First, we used BEDTools (Quinlan & Hall, 2010) to quantify the coverage of each predicted peak using input (mock) and immunoprecipitation (IP, antibody against RBP) BAM files. Next, for each peak, the relative information content was defined as p_i_ x log_2_ (p_i_/q_i_), where p and q are the sums of reads mapping to the peak in IP and negative control respectively. The information content was used to calculate the Irreproducible Discovery Rate (IDR) (Li et al., 2011), a parameter indicating reproducible peaks across biological replicates. A significant and reproducible peak was defined meeting an IDR cut-off < 0.01, p-value ≤ 10^−5^ and fold-enrichment (FC) > 8. In case two or more peaks overlapped the same genomic region, the most significant one was included in the peak table. Additionally, non-overlapping peaks were pooled into a single table, in order to get a complete set in both cell lines. While CLIP data was produced in a non-cardiac setting, CLIP signals are usually preserved among similarly expressed genes of the same RBP independent of the cell line, with peak differences instead reflecting cell type-specific expression rather than binding affinity (Van Nostrand et al., 2020). Additionally, for the muscle-specific splicing repressor RBM20, which was not part of the ENCODE dataset but included for its importance for cardiac splicing and heart disease (Maatz et al., 2014; Guo et al., 2012), significant rat RBM20 HITS-CLIP targets were obtained from Maatz et al. (Maatz et al., 2014) and converted to GRCh38.p10/hg38 genomic coordinates. Only 143 RBPs with expression in human heart tissue were kept (mean FPKM across samples >1; 142 ENCODE RBPs and RBM20). Overall, we retrieved an average of 4,300 eCLIP-seq peaks per experiment. Finally, we mapped these peaks to the annotated transcriptome (Ensembl v.87) and, for each RBP experiment, all the genes supported by at least one CLIP-seq peak were defined as putative target genes.

### RBP-target correlation and clustering

For RBP-target correlations and clustering we included genes expressed in the human heart (mean FPKM across samples > 1) with at least one Ribo-seq and mRNA-seq read in a minimum of 20 samples (n = 11,387). Next, for pairwise complete observations, we calculated Spearman correlations between the expression level of the RBP (as measured by Ribo-seq) and either target gene mRNA-seq counts or translational efficiency. Only target genes that showed a significant (p_adj_ ≤ 0.05) correlation after correction for multiple testing using the Benjamini-Hochberg approach (Benjamini & Hochberg, 1995) were retained for downstream analyses. The computed RBP-target correlation matrix was used to calculate the Euclidean distance followed by hierarchical clustering, in order to group RBPs with similar consequences on their target genes. Cluster visualization was done using heatmap.3 (https://github.com/obigriffith/biostar-tutorials/tree/master/Heatmaps).

### Target gene enrichment

To identify RBPs that are putative modulators of target gene mRNA abundance and/or TE, we calculated the frequency with which target genes supported with CLIP-seq data correlated significantly with each RBP. We leveraged the significance of these correlating associations by generating 100,000 equally sized sets of theoretical targets out of all translated genes in the human heart; an approach that has previously been shown to be highly effective (Chothani et al., 2019). For each set, we quantified the amount of significantly correlating genes and compared the theoretical distribution against the actual observation applying an empirical test:

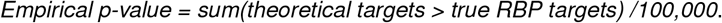

Empirical p-values were corrected for multiple testing (Benjamini-Hochberg method). RBPs that showed a significant (p_adj_ ≤ 0.05) enrichment of correlating CLIP-derived target genes were considered as putative regulators of mRNA abundance (n = 58) and/or TE (n = 37). It should be noted that, because of the fixed number of generated random sets, the minimum empirical p-value that can be calculated after correction for multiple testing is 5.25 × 10^−5^. Hence, the empirical test cannot quantify the strength of significance for a specific observation. Instead, we calculated Glass’ Delta (Δ)(Freeman et al., 1986) as a measure of the effect size, which is defined as the difference between the two target sets divided by the standard deviation of the theoretical group.

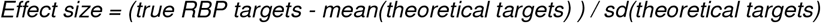

### RBP expression across GTEx tissues

To determine the patterns of expression of each RBP across human tissues, we obtained expression data from the Genotype-Tissue Expression (GTEx) Project (GTEx Consortium, 2013), a database that comprises a large set of samples corresponding to 54 human tissues. We used these data to determine the number of tissues with detectable (average TPM ≥1) or high (average TPM ≥10) expression of a given RBP. An RBP was categorized as ubiquitously expressed if expression was detected in more than 30 tissues.

### Replication of target regulation using a public fibroblast cohort

We retrieved raw RNA-seq and Ribo-seq data from a cohort of 20 primary cardiac fibroblast cultures stimulated with TGF-beta (Chothani et al., 2019) and used it as a replication cohort. Raw data are available via the gene expression omnibus (GEO submission: GSE131112, GSE123018, GSE131111) repository. Read pre-processing, mapping, gene quantification and correlation analysis were done following the same procedures described above for the heart datasets (see ‘Read mapping and gene quantification’ and ‘RBP-target correlation and clustering’ subsections). To prove that the regulatory effect of RBPs in target translational regulation can be replicated in an independent dataset, we quantified the fraction of RBP-target correlations with similar direction of regulation in both fibroblast and human heart cohorts. Statistical significance of the observed replications was evaluated by running 10,000 permutations of the correlation coefficients in fibroblasts and comparing the fraction of shared directionality between both cohorts in observed and randomized sets.

### Analysis of differential exon splicing

To evaluate whether RBM20 could influence the TE of target genes by modulating isoform production ratios (exon in- or exclusion), we estimated exon splicing rates by calculating the percentage spliced in (PSI) for all exons of known and correlating RBM20 target genes, as described previously (Schafer et al., 2015a). For PSI calculation, we re-mapped the 80 paired-end cardiac mRNA-seq (2 × 101nt) datasets to improve splice site coverage using STAR v2.5.2b (Dobin et al., 2013), allowing a maximum of 6 mismatches.

### Functional analysis of RBP associations and target genes

Known and predicted RBP-RBP interactions were retrieved from the STRING database (Szklarczyk et al., 2019) with confidence network edges and default settings. Moreover, we assigned biological functions to define gene targets with gProfiler2 v0.1.9 (archive revision fof4439, (Raudvere et al., 2019)) and extracted enriched sets of ‘child’ and ‘parent’ GO terms for the individual sets mRNA and TE targets (p_adj_ ≤ 0.05). Significant GO terms that involve a minimum of20 and a maximum of 500 genes were considered, to avoid the inclusion of too general terms that show significance due to beneficial input to term size ratios.

### Analysis of minimum free energy in 5’ UTRs

We predicted 5’ UTR secondary structures through energy minimization using RNAfold from the Vienna Package v2.4 (Lorenz et al., 2011). Using the 5’ UTR sequence of each target gene as input, minimum free energies (MFE) were calculated and length-normalized to observe differences in UTR complexity for target genes that are positively or negatively correlating with RBPs.

### General remarks on statistical analysis

Statistical analysis and generation of figures was done using R v3.6.2 (R Core Team (2020), 2020). A full list of tools and methods used for data analysis is stated in each corresponding Methods section. Statistical parameters such as n, median/mean, standard deviation (SD) and significance are named in the figures and/or the figure legends. The “p_adj_” indicates the significance after correction for multiple testing using the Benjamini-Hochberg approach (Benjamini & Hochberg, 1995). The “*n*” represents the number of RBPs in **Figure S3A**.

## SUPPLEMENTARY TABLES

**Table S1: Analysis information for 143 RBPs** Table with all 143 cardiac expressed RBPs, number of mRNA and TE correlating targets and significance of correlations, clusters of coregulated RBPs, and average RNA expression levels in human left ventricle.

**Table S2: Multifunctional RBPs** Table with all 21 multifunctional RBPs, number of target genes per molecular trait, and names and significance of the best 5 GO enrichment results.

**Table S3: Multifunctional RBPs’ localization** Table with all 21 multifunctional RBPs and their cellular localization (0: absent; 1: present).

**Figure S1.**
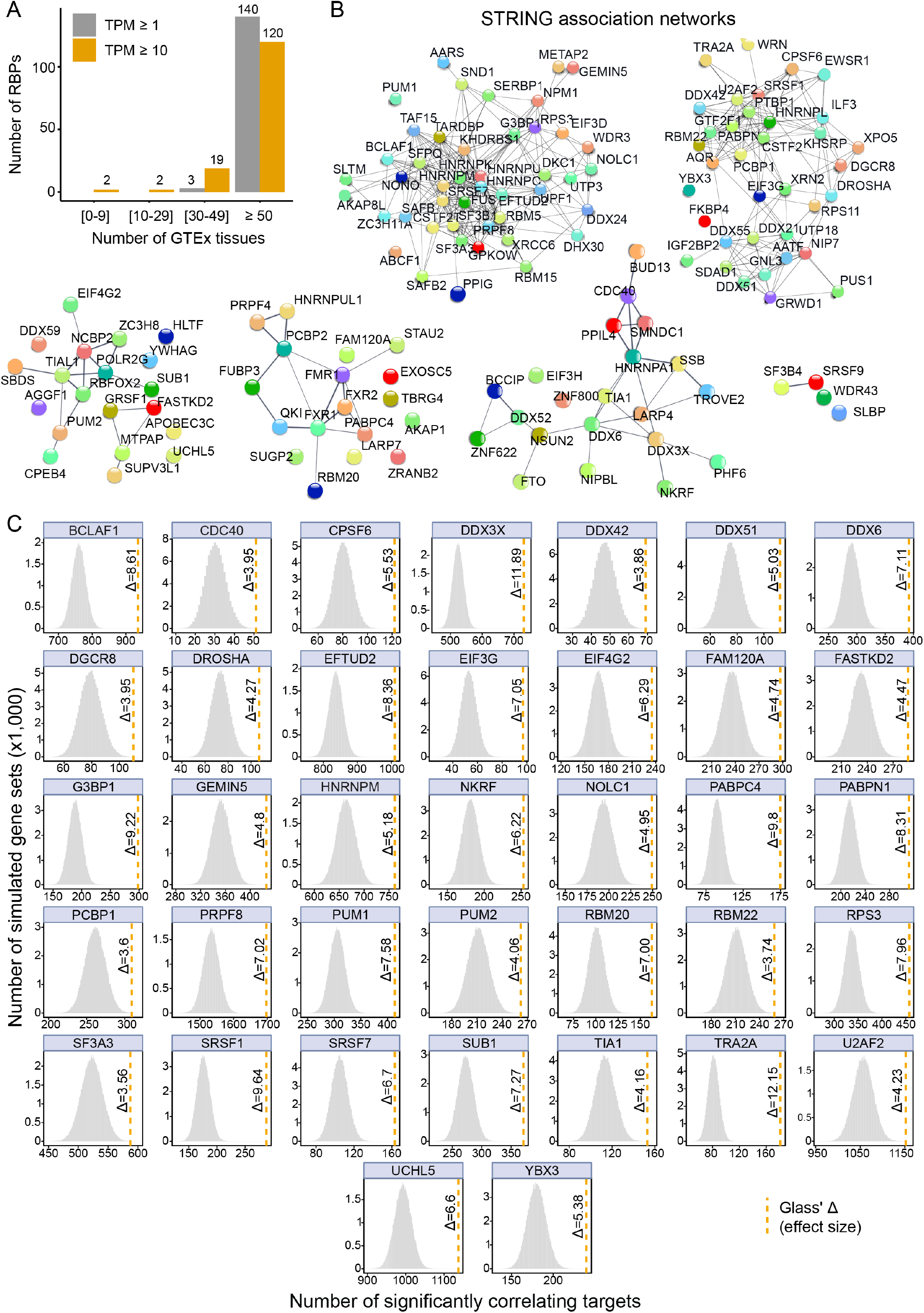
RNA-binding protein abundance predicts target translational regulation. **(A)** Bar plot displaying the patterns of expression of the 143 RBPs across tissues. Average expression values in transcript per million (TPM) units were retrieved from the Genotype-Tissue Expression (GTEx) Project. Most of the RBPs are ubiquitously expressed across human tissues. **(B)** STRING protein-protein association networks from six coregulated RBP clusters (see also **Figure 1A**). Most of the clustered RBPs are involved in known functional interactions. **(C)** Heatmaps with Glass’ Δ scores for all 37 TE-RBPs quantifying the effect size of the witness effects for significant TE correlations.

**Figure S2.**
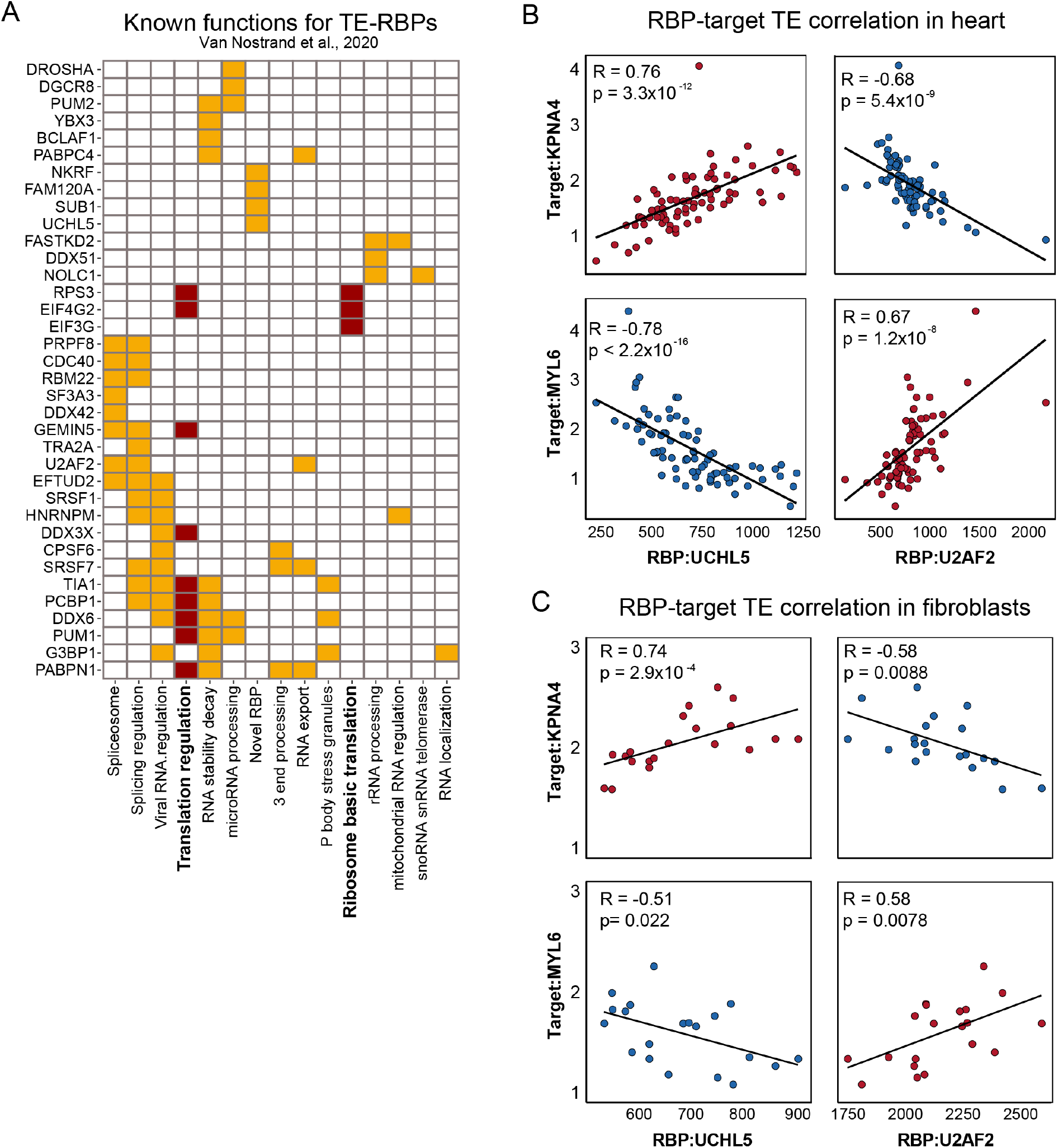
CLIP analysis identifies coregulated *in vivo* targets and novel master translational regulators in the human heart. **(A)** Described functions by Van Nostrand et al. (Van Nostrand et al., 2020) for the set of TE-RBPs. Functions related to translation (translation regulation and ribosome basic translation) are highlighted with dark red boxes. **(B-C)** Scatter plots representing the correlation of heart **(B)** and primary cardiac fibroblasts **(C)** translational efficiencies between UCHL5 and U2AF2 and two shared targets, KPNA4 and MYL6. UCHL5 and U2AF2 have marked opposite effects on their shared targets, indicative of a competitive effect replicated in two independent datasets. Scores and level of significance of the two Spearman’s correlations are displayed.

**Figure S3.**
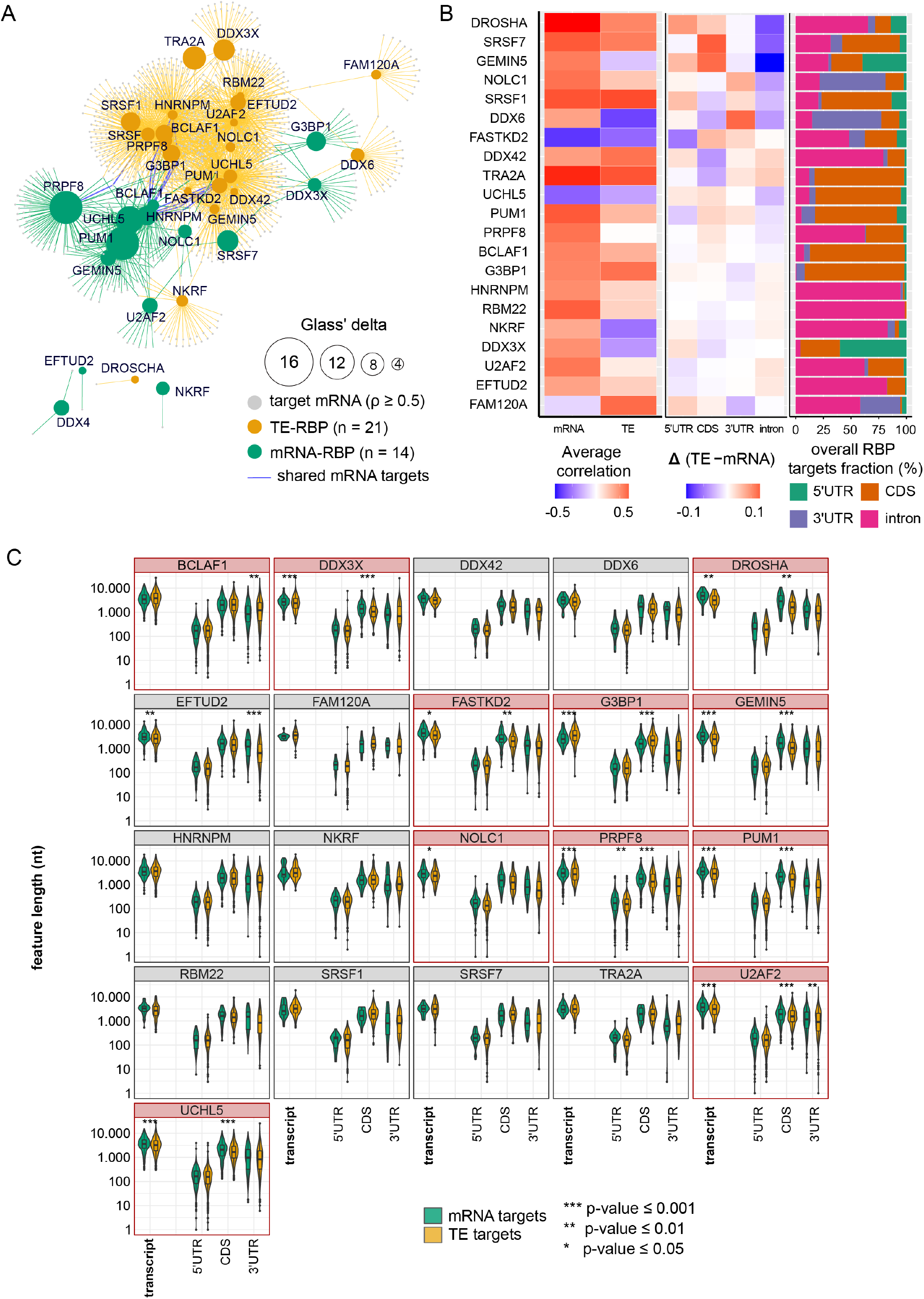
Multifunctional RBPs regulate translation of distinct sets of target genes. **(A)** Network representing multifunctional RBP-target interactions for both mRNA-RBPs (green) and TE-RBPs (brown) of strong correlating pairs. Blue lines indicate shared targets in both mRNA abundance and TE regulation of the same RBP. **(B)** Left: heatmap representing the average mRNA and TE RBP-target correlation values for all 21 multifunctional RBPs. Middle: heatmap representing differences in the relative proportion of feature binding sites (TE-mRNA) for all 21 multifunctional RBPs. Right: bar plot showing the overall proportion of feature binding sites for all 21 multifunctional RBPs. **(C)** Box plots with 5’ UTR, CDS, and 3’ UTR sequence lengths in nucleotides for mRNA and TE targets corresponding to the set of 21 multifunctional RBPs. For each target gene, the most abundant isoform is represented.

## REFERENCES

Anders S., Pyl P.T. & Huber W. (2015). HTSeq–a Python framework to work with high-throughput sequencing data. Bioinformatics (Oxford, England) 31.(2), 166–9.

Backlund M., Stein F., Rettel M., Schwarzl T., Perez-Perri J. I., Brosig A., Zhou Y., Neu-Yilik G., Hentze M.W. & Kulozik A. E. (2020). Plasticity of nuclear and cytoplasmic stress responses of RNA-binding proteins. Nucleic acids research 48.(9), 4725–4740.

Beckmann B. M., Castello A. & Medenbach J. (2016). The expanding universe of ribonucleoproteins: of novel RNA-binding proteins and unconventional interactions. Pflugers Archiv : European journal of physiology 468.(6), 1029–40.

Benjamini Y. & Hochberg Y. (1995). Controlling the False Discovery Rate: A Practical and Powerful Approach to Multiple Testing. Journal of the Royal Statistical Society: Series B (Methodological) 57.(1), 289–300.

Buchan J. R. (2014). mRNP granules. Assembly, function, and connections with disease. RNA biology 11.(8), 1019–30.

Burgess H. M., Richardson W. A., Anderson R. C., Salaun C., Graham S.V. & Gray N. K. (2011). Nuclear relocalisation of cytoplasmic poly(A)-binding proteins PABP1 and PABP4 in response to UV irradiation reveals mRNA-dependent export of metazoan PABPs. Journal of cell science 124.(Pt 19), 3344–55.

Calviello L., Venkataramanan S., Rogowski K. J., Wyler E., Wilkins K., Tejura M., Thai B., Krol J., Filipowicz W., Landthaler M., et al. (2020). DDX3 depletion represses translation of mRNAs with complex 5’ UTRs. bioRxiv, 589218.

Castello A., Fischer B., Frese C. K., Horos R., Alleaume A.-M., Foehr S., Curk T., Krijgsveld J. & Hentze M. W. (2016). Comprehensive Identification of RNA-Binding Domains in Human Cells. Molecular cell 63.(4), 696–710.

Caudron-Herger M., Rusin S. F., Adamo M. E., Seiler J., Schmid V. K., Barreau E., Kettenbach A.N. & Diederichs S. (2019). R-DeeP: Proteome-wide and Quantitative Identification of RNA-Dependent Proteins by Density Gradient Ultracentrifugation. Molecular cell 75.(1), 184–199.e10.

Chen T.-M., Lai M.-C., Li Y.-H., Chan Y.-L., Wu C.-H., Wang Y.-M., Chien C.-W., Huang S.-Y., Sun H.S. & Tsai S.-J. (2019). hnRNPM induces translation switch under hypoxia to promote colon cancer development. EBioMedicine 41, 299–309.

Cheng S., Fahmi N. A., Park M., Chang J.-W., Sun J., Thao K., Yeh H.-S., Zhang W. & Yong J. (2020). mTOR-driven widespread exon skipping renders multifaceted gene regulation and proteome complexity. bioRxiv, 2020.02.27.967737.

Chothani S., Schäfer S., Adami E., Viswanathan S., Widjaja A. A., Langley S. R., Tan J., Wang M., Quaife N. M., Jian Pua C., et al. (2019). Widespread Translational Control of Fibrosis in the Human Heart by RNA-Binding Proteins. Circulation 140.(11), 937–951.

Cirillo L., Cieren A., Barbieri S., Khong A., Schwager F., Parker R. & Gotta M. (2020). UBAP2L Forms Distinct Cores that Act in Nucleating Stress Granules Upstream of G3BP1. Current biology : CB 30.(4), 698–707.e6.

Copsey A. C., Cooper S., Parker R., Lineham E., Lapworth C., Jallad D., Sweet S. & Morley S. J. (2017). The helicase, DDX3X, interacts with poly(A)-binding protein 1 (PABP1) and caprin-1 at the leading edge of migrating fibroblasts and is required for efficient cell spreading. The Biochemical journal 474.(18), 3109–3120.

Costa M., Ochem A., Staub A. & Falaschi A. (1999). Human DNA helicase VIII: a DNA and RNA helicase corresponding to the G3BP protein, an element of the ras transduction pathway. Nucleic acids research 27.(3), 817–21.

Damianov A., Ying Y., Lin C.-H., Lee J.-A., Tran D., Vashisht A. A., Bahrami-Samani E., Xing Y., Martin K. C., Wohlschlegel J. A., et al. (2016). Rbfox Proteins Regulate Splicing as Part of a Large Multiprotein Complex LASR. Cell 165.(3), 606–19.

Dassi E. (2017). Handshakes and Fights: The Regulatory Interplay of RNA-Binding Proteins. Frontiers in molecular biosciences 4, 67.

Davis C. A., Hitz B. C., Sloan C. A., Chan E. T., Davidson J. M., Gabdank I., Hilton J. A., Jain K., Baymuradov U. K., Narayanan A. K., et al. (2018). The Encyclopedia of DNA elements (ENCODE): data portal update. Nucleic acids research 46.(D1), D794–D801.

De Gaudenzi J. G., Jäger A. V., Izcovich R. & Campo V. A. (2016). Insights into the Regulation of mRNA Processing of Polycistronic Transcripts Mediated by DRBD4/PTB2, a Trypanosome Homolog of the Polypyrimidine Tract-Binding Protein. The Journal of eukaryotic microbiology.

Díaz-Muñoz M.D. & Turner M. (2018). Uncovering the Role of RNA-Binding Proteins in Gene Expression in the Immune System. Frontiers in immunology 9.(MAY), 1094.

Dobin A., Davis C. A., Schlesinger F., Drenkow J., Zaleski C., Jha S., Batut P., Chaisson M. & Gingeras T. R. (2013). STAR: ultrafast universal RNA-seq aligner. Bioinformatics (Oxford, England) 29.(1), 15–21.

Dong G., Liang F., Sun B., Wang C., Liu Y., Guan X., Yang B., Xiu C., Yang N., Liu F., et al. (2019). Presence and function of stress granules in atrial fibrillation. PLoS ONE 14.(4).

Drummond D.A. & Wilke C. O. (2008). Mistranslation-induced protein misfolding as a dominant constraint on coding-sequence evolution. Cell 134.(2), 341–52.

Fischer J. W., Busa V. F., Shao Y. & Leung A. K. L. (2020). Structure-Mediated RNA Decay by UPF1 and G3BP1. Molecular cell 78.(1), 70–84.e6.

Freeman P. R., Hedges L.V. & Olkin I. (1986). Statistical Methods for Meta-Analysis. Biometrics 42.(2), 454.

Gerstberger S., Hafner M. & Tuschl T. (2014). A census of human RNA-binding proteins. Nature reviews. Genetics 15.(12), 829–45.

Gilbertson S., Federspiel J. D., Hartenian E., Cristea I.M. & Glaunsinger B. (2018). Changes in mRNA abundance drive shuttling of RNA binding proteins, linking cytoplasmic RNA degradation to transcription. eLife 7.

Gray N. K., Hrabálková L., Scanlon J.P. & Smith R. W. P. (2015). Poly(A)-binding proteins and mRNA localization: who rules the roost? Biochemical Society transactions 43.(6), 1277–84.

GTEx Consortium (2013). The Genotype-Tissue Expression (GTEx) project. Nature genetics 45.(6), 580–5.

Guenther U.-P., Weinberg D. E., Zubradt M. M., Tedeschi F. A., Stawicki B. N., Zagore L. L., Brar G. A., Licatalosi D. D., Bartel D. P., Weissman J. S., et al. (2018). The helicase Ded1p controls use of near-cognate translation initiation codons in 5’ UTRs. Nature 559.(7712), 130–134.

Guo W., Schafer S., Greaser M. L., Radke M. H., Liss M., Govindarajan T., Maatz H., Schulz H., Li S., Parrish A. M., et al. (2012). RBM20, a gene for hereditary cardiomyopathy, regulates titin splicing. Nature medicine 18.(5), 766–73.

Hannon G. (2010). FASTX-Toolkit.

Harvey R. F., Smith T. S., Mulroney T., Queiroz R. M. L., Pizzinga M., Dezi V., Villenueva E., Ramakrishna M., Lilley K.S. & Willis A. E. (2018a). Trans-acting translational regulatory RNA binding proteins. Wiley interdisciplinary reviews. RNA 9.(3), e1465.

Harvey S. E., Xu Y., Lin X., Gao X. D., Qiu Y., Ahn J., Xiao X. & Cheng C. (2018b). Coregulation of alternative splicing by hnRNPM and ESRP1 during EMT. RNA (New York, N.Y.) 24.(10), 1326–1338.

Hilliker A., Gao Z., Jankowsky E. & Parker R. (2011). The DEAD-box protein Ded1 modulates translation by the formation and resolution of an eIF4F-mRNA complex. Molecular cell 43.(6), 962–72.

Hinnebusch A. G. (2014). The scanning mechanism of eukaryotic translation initiation. Annual review of biochemistry 83, 779–812.

Katahira J. (2015). Nuclear export of messenger RNA. Genes 6.(2), 163–84.

Kim J., Park R. Y., Chen J.-K., Kim J., Jeong S. & Ohn T. (2014). Splicing factor SRSF3 represses the translation of programmed cell death 4 mRNA by associating with the 5’-UTR region. Cell death and differentiation 21.(3), 481–90.

Kim Y. & Myong S. (2016). RNA Remodeling Activity of DEAD Box Proteins Tuned by Protein Concentration, RNA Length, and ATP. Molecular cell 63.(5), 865–76.

Kong X., Chen Y.-Y., Lin J., Flowers E., Van Nostrand E., Blue S. M., Chau J., Ma C. I.-H., Mohr I., Thai R., et al. (2019). The cohesin loader NIPBL interacts with pre-ribosomal RNA and treacle to regulate ribosomal RNA synthesis. bioRxiv.

Laver J. D., Ly J., Winn A. K., Karaiskakis A., Lin S., Nie K., Benic G., Jaberi-Lashkari N., Cao W. X., Khademi A., et al. (2020). The RNA-Binding Protein Rasputin/G3BP Enhances the Stability and Translation of Its Target mRNAs. Cell reports 30.(10), 3353–3367.e7.

Leppek K., Das R. & Barna M. (2018). Functional 5’ UTR mRNA structures in eukaryotic translation regulation and how to find them. Nature reviews. Molecular cell biology 19.(3), 158–174.

Lewis B. P., Green R.E. & Brenner S. E. (2003). Evidence for the widespread coupling of alternative splicing and nonsense-mediated mRNA decay in humans. Proceedings of the National Academy of Sciences of the United States of America 100.(1), 189–192.

Li Q., Brown J. B., Huang H. & Bickel P. J. (2011). Measuring reproducibility of high-throughput experiments. The Annals of Applied Statistics 5.(3), 1752–1779.

Liberman N., Marash L. & Kimchi A. (2009). The translation initiation factor DAP5 is a regulator of cell survival during mitosis. Cell cycle (Georgetown, Tex.) 8.(2), 204–9.

Lim C. S.T, Wardell S. J., Kleffmann T. & Brown C. M. (2018). The exon-intron gene structure upstream of the initiation codon predicts translation efficiency. Nucleic acids research 46.(9), 4575–4591.

Lorenz R., Bernhart S. H., Höner Zu Siederdissen C., Tafer H., Flamm C., Stadler P.F. & Hofacker I. L. (2011). ViennaRNA Package 2.0. Algorithms for molecular biology : AMB 6.(1), 26.

Love M. I., Huber W. & Anders S. (2014). Moderated estimation of fold change and dispersion for RNA-seq data with DESeq2. Genome biology 15.(12), 550.

Luo E.-C., Nathanson J. L., Tan F. E., Schwartz J. L., Schmok J. C., Shankar A., Markmiller S., Yee B. A., Sathe S., Pratt G. A., et al. (2020). Large-scale tethered function assays identify factors that regulate mRNA stability and translation. Nature structural & molecular biology 27.(10), 989–1000.

Maatz H., Jens M., Liss M., Schafer S., Heinig M., Kirchner M., Adami E., Rintisch C., Dauksaite V., Radke M. H., et al. (2014). RNA-binding protein RBM20 represses splicing to orchestrate cardiac pre-mRNA processing. The Journal of clinical investigation 124.(8), 3419–30.

Malinová A., Cvačková Z., Matějů D., Hořejší Z., Abéza C., Vandermoere F., Bertrand E., Staněk D. & Verheggen C. (2017). Assembly of the U5 snRNP component PRPF8 is controlled by the HSP90/R2TP chaperones. The Journal of cell biology 216.(6), 1579–1596.

Mallam A. L., Sae-Lee W., Schaub J. M., Tu F., Battenhouse A., Jang Y. J., Kim J., Wallingford J. B., Finkelstein I. J., Marcotte E. M., et al. (2019). Systematic Discovery of Endogenous Human Ribonucleoprotein Complexes. Cell reports 29.(5), 1351–1368.e5.

Maslon M. M., Heras S. R., Bellora N., Eyras E. & Cáceres J. F. (2014). The translational landscape of the splicing factor SRSF1 and its role in mitosis. eLife 3, e02028.

Matsuki H., Takahashi M., Higuchi M., Makokha G. N., Oie M. & Fujii M. (2013). Both G3BP1 and G3BP2 contribute to stress granule formation. Genes to cells : devoted to molecular & cellular mechanisms 18.(2), 135–46.

Mauger D. M., Cabral B. J., Presnyak V., Su S. V., Reid D. W., Goodman B., Link K., Khatwani N., Reynders J., Moore M. J., et al. (2019). mRNA structure regulates protein expression through changes in functional half-life. Proceedings of the National Academy of Sciences of the United States of America 116.(48), 24075–24083.

Mo J., Liang H., Su C., Li P., Chen J. & Zhang B. (2021). DDX3X: structure, physiologic functions and cancer. Molecular cancer 20.(1), 38.

Müller-McNicoll M. & Neugebauer K. M. (2013). How cells get the message: dynamic assembly and function of mRNA-protein complexes. Nature reviews. Genetics 14.(4), 275–87.

Nott A., Le Hir H. & Moore M. J. (2004). Splicing enhances translation in mammalian cells: an additional function of the exon junction complex. Genes & development 18.(2), 210–22.

Pelletier J. & Sonenberg N. (1985). Insertion mutagenesis to increase secondary structure within the 5’ noncoding region of a eukaryotic mRNA reduces translational efficiency. Cell 40.(3), 515–26.

Qian W., Yang J.-R., Pearson N. M., Maclean C. & Zhang J. (2012). Balanced codon usage optimizes eukaryotic translational efficiency. PLoS genetics 8.(3), e1002603.

Quinlan A.R. & Hall I. M. (2010). BEDTools: a flexible suite of utilities for comparing genomic features. eng. Bioinformatics (Oxford, England) 26.(6), 841–2.

R Core Team (2020) (2020). R: A language and environment for statistical computing.

Raudvere U., Kolberg L., Kuzmin I., Arak T., Adler P., Peterson H. & Vilo J. (2019). g:Profiler: a web server for functional enrichment analysis and conversions of gene lists (2019 update). Nucleic acids research 47.(W1), W191–W198.

Ray D., Kazan H., Cook K. B., Weirauch M. T., Najafabadi H. S., Li X., Gueroussov S., Albu M., Zheng H., Yang A., et al. (2013). A compendium of RNA-binding motifs for decoding gene regulation. Nature 499.(7457), 172–7.

Sahoo P. K., Lee S. J., Jaiswal P. B., Alber S., Kar A. N., Miller-Randolph S., Taylor E. E., Smith T., Singh B., Ho T. S.-Y., et al. (2018). Axonal G3BP1 stress granule protein limits axonal mRNA translation and nerve regeneration. Nature communications 9.(1), 3358.

Schafer S., Miao K., Benson C. C., Heinig M., Cook S.A. & Hubner N. (2015a). Alternative Splicing Signatures in RNA-seq Data: Percent Spliced in (PSI). Current protocols in human genetics 87, 11.16.1–11.16.14.

Schafer S., Adami E., Heinig M., Rodrigues K. E. C., Kreuchwig F., Silhavy J., Heesch S. van, Simaite D., Rajewsky N., Cuppen E., et al. (2015b). Translational regulation shapes the molecular landscape of complex disease phenotypes. Nature communications 6, 7200.

Sen N. D., Zhou F., Ingolia N.T. & Hinnebusch A. G. (2015). Genome-wide analysis of translational efficiency reveals distinct but overlapping functions of yeast DEAD-box RNA helicases Ded1 and eIF4A. Genome research 25.(8), 1196–205.

Sharma D., Zagore L. L., Brister M. M., Ye X., Crespo-Hernández C. E., Licatalosi D.D. & Jankowsky E. (2021). The kinetic landscape of an RNA-binding protein in cells. Nature 591.(7848), 152–156.

Shen L. & Pelletier J. (2020). General and Target-Specific DExD/H RNA Helicases in Eukaryotic Translation Initiation. International journal of molecular sciences 21.(12).

Shi Z., Fujii K., Kovary K. M., Genuth N. R., Röst H.L., Teruel M.N. & Barna M. (2017). Heterogeneous Ribosomes Preferentially Translate Distinct Subpools of mRNAs Genome-wide. Molecular cell 67.(1), 71–83.e7.

Simsek D., Tiu G. C., Flynn R. A., Byeon G. W., Leppek K., Xu A. F., Chang H.Y. & Barna M. (2017). The Mammalian Ribo-interactome Reveals Ribosome Functional Diversity and Heterogeneity. Cell 169.(6), 1051–1065.e18.

Soto-Rifo R., Rubilar P. S., Limousin T., Breyne S. de, Décimo D. & Ohlmann T. (2012). DEAD-box protein DDX3 associates with eIF4F to promote translation of selected mRNAs. The EMBO journal 31.(18), 3745–56.

Sutherland L. C., Wang K. & Robinson A. G. (2010). RBM5 as a putative tumor suppressor gene for lung cancer.

Szklarczyk D., Gable A. L., Lyon D., Junge A., Wyder S., Huerta-Cepas J., Simonovic M., Doncheva N. T., Morris J. H., Bork P., et al. (2019). STRING v11: protein-protein association networks with increased coverage, supporting functional discovery in genome-wide experimental datasets. Nucleic acids research 47.(D1), D607–D613.

Trendel J., Schwarzl T., Horos R., Prakash A., Bateman A., Hentze M.W. & Krijgsveld J. (2019). The Human RNA-Binding Proteome and Its Dynamics during Translational Arrest. Cell 176.(1-2), 391–403.e19.

Van Heesch S., Witte F., Schneider-Lunitz V., Schulz J. F., Adami E., Faber A. B., Kirchner M., Maatz H., Blachut S., Sandmann C.-L., et al. (2019). The Translational Landscape of the Human Heart. Cell 178.(1), 242–260.e29.

Van Nostrand E. L., Freese P., Pratt G. A., Wang X., Wei X., Xiao R., Blue S. M., Chen J.-Y., Cody N. A. L., Dominguez D., et al. (2020). A large-scale binding and functional map of human RNA-binding proteins. Nature 583.(7818), 711–719.

Wang S. E., Brooks A. E. S., Poole A.M. & Simoes-Barbosa A. (2020). Determinants of translation efficiency in the evolutionarily-divergent protist Trichomonas vaginalis. BMC molecular and cell biology 21.(1), 54.

Weber R, Kleemann L, Hirschberg I, Chung M, Valkov E I. C. (2021). DAP5 enables translation re-initiation on structured messenger RNAs. bioRxiv.

Yang W., Ru Y., Ren J., Bai J., Wei J., Fu S., Liu X., Li D. & Zheng H. (2019). G3BP1 inhibits RNA virus replication by positively regulating RIG-I-mediated cellular antiviral response. Cell death & disease 10.(12), 946.

Yao T., Song L., Jin J., Cai Y., Takahashi H., Swanson S. K., Washburn M. P., Florens L., Conaway R. C., Cohen R. E., et al. (2008). Distinct modes of regulation of the Uch37 deubiquitinating enzyme in the proteasome and in the Ino80 chromatin-remodeling complex. Molecular cell 31.(6), 909–17.

Yedavalli V. S.R. K., Neuveut C., Chi Y.-H., Kleiman L. & Jeang K.-T. (2004). Requirement of DDX3 DEAD box RNA helicase for HIV-1 Rev-RRE export function. Cell 119.(3), 381–92.

Ying S. & Khaperskyy D. A. (2021). UV damage induces G3BP1-dependent stress granule formation that is not driven by mTOR inhibition-mediated translation arrest. Journal of Cell Science.

Yugami M., Okano H., Nakanishi A. & Yano M. (2020). Analysis of the nucleocytoplasmic shuttling RNA-binding protein HNRNPU using optimized HITS-CLIP method. PloS one 15.(4), e0231450.

Zhong Y., Karaletsos T., Drewe P., Sreedharan V. T., Kuo D., Singh K., Wendel H.-G. & Rätsch G. (2017). RiboDiff: detecting changes of mRNA translation efficiency from ribosome footprints. Bioinformatics (Oxford, England) 33.(1), 139–141.

Zhou Z., Dang Y., Zhou M., Li L., Yu C.-H., Fu J., Chen S. & Liu Y. (2016). Codon usage is an important determinant of gene expression levels largely through its effects on transcription. Proceedings of the National Academy of Sciences of the United States of America 113.(41), E6117–E6125.

